# Virulence and genomic analysis of *Puccinia coronata* f. sp. *avenae* in Australia identifies new races and a new lineage in 2024

**DOI:** 10.1101/2025.11.08.687394

**Authors:** Zhouyang Su, David Lewis, Eva C. Henningsen, Duong T. Nguyen, Jana Sperschneider, Peter N. Dodds, Melania Figueroa

## Abstract

Crown rust, caused by *Puccinia coronata* f. sp. *avenae* (*Pca*), remains a persistent threat to oat production in Australia. To monitor recent shifts in virulence and population structure, 30 *Pca* isolates collected during the 2024 growing season across major Australian oat-producing regions were analysed. Virulence analysis of 30 isolates using 52 oat differential lines identified 25 unique races that were not detected in previous years. Whole-genome sequencing of 28 of these isolates were analysed in the context of a broader historical Australian and international genomic datasets including isolates from Taiwan, South Africa, USA. Results confirmed the uniqueness of the Australian *Pca* population and revealed well-established genotypic lineages persisting over multiple years, with L18 and L16 being dominant. Notably, L16 was again present in Western Australia after being undetected in 2023, while L18 maintained its prevalence for a third consecutive season. Beyond these dominant groups, phylogenetic analysis and a *k*-mer containment analysis also identified a novel and genetically distinct lineage, designated as L19, represented by one isolate collected in WA. To add to the characterisation of lineage L19, we recorded virulence phenotypes on a small collection of current commercial cultivars. These findings enhance understanding of *Pca* diversity and emphasise the importance of surveillance approaches that integrate phenotypic and genomic surveillance.

## Manuscript body

Oat crown rust (OCR), caused by *Puccinia coronata* f. sp. *avenae* (*Pca*), is the most destructive foliar disease for oat production worldwide (Nazareno et al. 2018). The disease typically causes yield losses of 10% - 40% under epidemic conditions (Nazareno et al. 2018). In a severe outbreak, yield losses have reached as high as 50% (USDA-ARS CDL 2014). In addition to reducing yield, the disease also affects grain quality and fodder value (Simons 1985), posing persistent challenges to both milling and forage oat production in Australia and other countries. While breeding for host resistance has proven to be effective (McDonald and Linde 2002), the durability of resistance genes is impacted by evolution of virulence in the *Pca* population (Carson 2011; McCallum et al. 2007; Moreau et al. 2024). Therefore, ongoing surveillance to detect changes in population structure and virulence profiles remains an essential component of OCR management strategies (Nazareno et al. 2018).

In Australia, routine surveys are conducted to monitor *Pca* virulence patterns and evaluate the effectiveness of *Pc* resistance genes (Henningsen et al. 2024b; Nguyen et al. 2025; Park et al. 2025). The high phenotypic diversity among *Pca* isolates is well-supported, with dozens of unique races identified each year (Henningsen et al. 2024b; Nguyen et al. 2025; Park et al. 2025). These findings underscore the dynamic nature of the *Pca* population. So far, a total of 18 genotypic lineages of *Pca* based on genome-wide variants have been characterised using 137 samples collected between 2020 and 2023 (Henningsen et al. 2024a). This framework enables a consistent lineage assignment and together with extensive haplotype resolved pangenome resources (Henningsen et al. 2024a) supports epidemiological inferences about migration and genetic recombination of *Pca* in Australia. Furthermore, a recent study highlights the need to complement phenotype-based assessments with genomic data as *Pca* isolates with similar virulence profiles can belong to genetically distinct lineages (Henningsen et al. 2024a). Therefore, full characterisation of the phenotypic and genotypic variability of *Pca* is critical to optimise genetic resistance deployment and understanding phenotypic shifts in the pathogen population that can impact the oat industry.

In this study, we report the virulence phenotypes of a collection of *Pca* isolates derived from 2024 and combine it with genotypic assignments based on whole genome sequencing. To bring these data into a global context, we leveraged published genomic data of historical and more recent international *Pca* populations (Henningsen et al. 2024a; Hewitt et al. 2024; Ho et al. 2025; Miller et al. 2020). Through a national network of plant pathologists and industry representatives, we obtained a total of 38 samples of *Pca* collected in 2024. These samples originated from New South Wales (*n*=7), Victoria (*n*=2), South Australia (*n*=5) and Western Australia (*n*=24) and were collected from wild oats (*n*=36), an unknown cultivated line (*n*=1), and an oat cultivar present in a field trial at Wagga Wagga, NSW. We note the lack of *Pca* sampling from Queensland, and low representation from New South Wales and Victoria, likely due to dry weather conditions (Bureau of Meteorology 2025, https://www.bom.gov.au/climate/maps/rainfall). Of the 38 *Pca* samples, 30 (NSW=6, VIC=2, SA=5 and WA=17) were revived after inoculation on the highly susceptible cultivars Swan and Marvelous following standard protocols (Miller et al. 2020). We proceeded with single pustule purification keeping purified *Pca* isolates for long term storage at -80°C (Henningsen et al. 2024a; Miller et al. 2020).

Virulence phenotypes of the 30 revived *Pca* isolates from the 2024 collection were determined using a panel of 52 differential lines as previously described (Henningsen et al. 2024b; Nguyen et al. 2025) (**Supplemental Table 1; Figure 1A**). A total of 25 unique races according to the 10-letter race coding system (Carson 2011; Chong et al. 2000; Nazareno et al. 2018) were identified (**Table 1**), confirming previous observations of the high diversity of virulence phenotypes for *Pca* in Australia. Two races were represented by more than one isolate: “BFBGMBDCG-” consisting of five isolates: 24WA12, 24WA13, 24WA14, 24WA22 and 24WA25, and “BJLQCCDCG-” consisting of two isolates: 24WA10 and 24WA23. All 25 races in 2024 represent new records according to previous collections (Henningsen et al. 2024b; Nguyen et al., 2025). We compared the distribution of virulence scores for the 2024 *Pca* isolates with those from previous years (Henningsen et al. 2024b; Nguyen et al. 2025) (**Supplemental Figure 1**). Overall, the collection from 2024 shows a shift toward less virulence than in the previous years 2022 and 2023 (*p*-value *<*0.001, Wilcoxon rank sum test) with 26 out of 52 oat differential lines showing less infection scores in 2024. The oat differential lines that were susceptible to the most isolates in this collection include Pc45 (*n*=29), Pc54 (*n*=29), Pc60 (*n*=26), Pc70 (*n*=25), and Pc46 (*n*=23). The oat lines Barcoo and Pc63 remained resistant to all isolates across these seasons, and observations from 2024 support low virulence for several oat lines such as Pc58, Pc59, Pc63 and WI X4361-9 mentioned in previous years (Henningsen et al. 2024b; Nguyen et al. 2025).

**Table 1.**
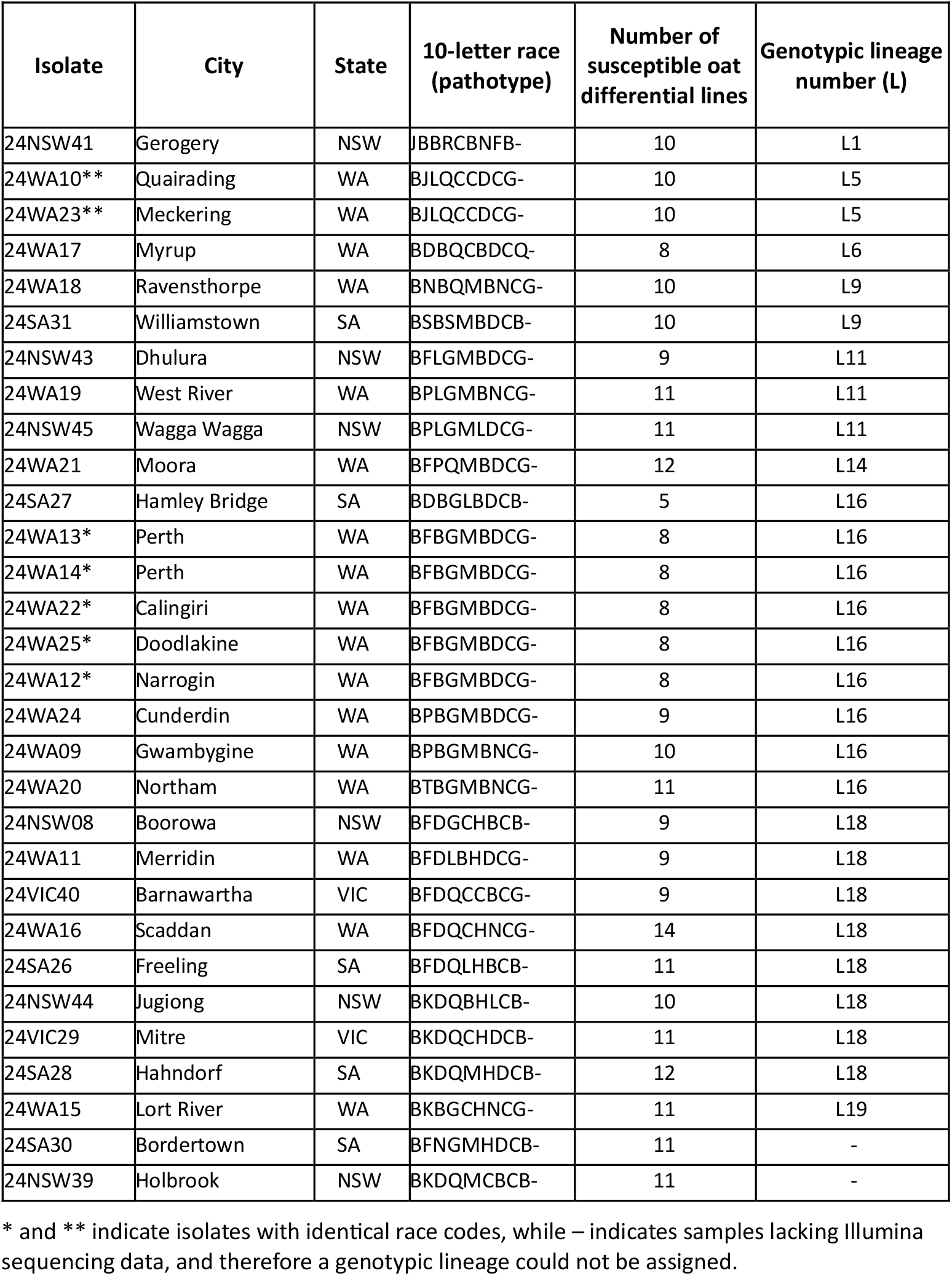
List of *Puccinia coronata* f. sp. *avenae* (*Pca*) isolates collected across Australia in 2024.

**Fig. 1.**
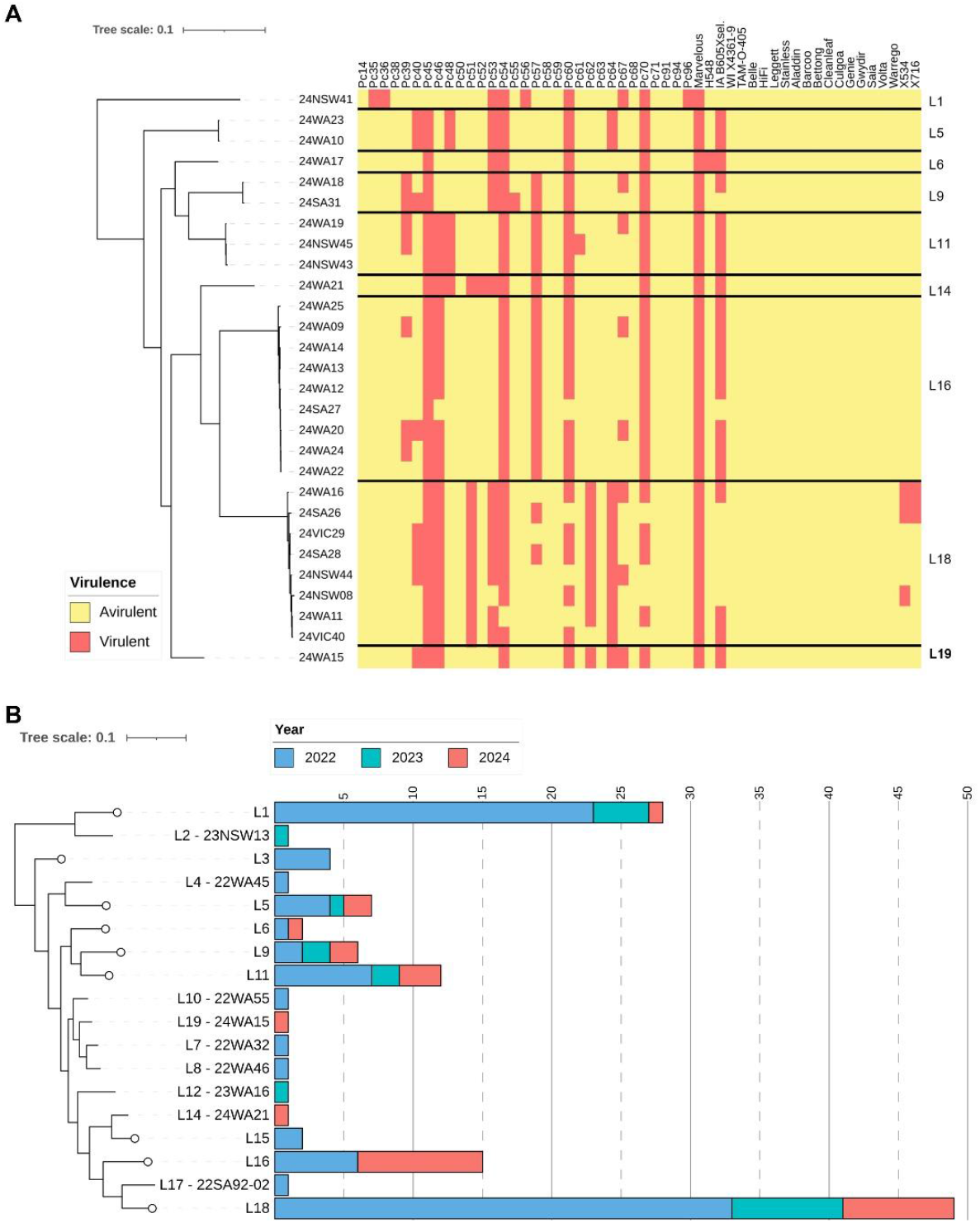
Australian *Pca* isolates collected in 2024. **A**. Heatmap showing virulence profiles of *Pca* isolates collected in 2024 (x-axis) on the oat differential lines (y-axis). High infection scores indicating high virulence (susceptibility) are shown in red, and lower infection scores indicating avirulence (resistance) are shown in yellow. WA = Western Australia, SA = South Australia, VIC = Victoria and NSW = New South Wales. **B**. Bar plots showing number of *Pca* isolates collected from 2022 to 2023 (Henningsen et al., 2024b; Nguyen et al., 2025) and this study that have been sequenced and assigned to genotypic lineages (Henningsen et al., 2024a).

To further explore population structure of the 2024 *Pca* isolates, whole-genome sequencing (WGS) data was generated. Genomic DNA was extracted for each isolate from 20–40 mg urediniospores using the OmniPrep Genomic DNA Isolation Kit (G-Biosciences) and sequencing libraries were prepared using the Illumina DNA PCR-Free Library Preparation Protocol. Sequencing on an Illumina NovaSeq X platform (150 bp paired-end reads, 15–30× coverage) was completed at the Biomolecular Resource Facility, Australian National University (Canberra, Australia). Two isolate samples (24SA30 and 24NSW39) did not pass WGS quality filters, and the remaining 28 samples were retained for the subsequent analysis (NSW=5, VIC=2, SA=4, WA=17) (**Supplemental Table 2)**. A Maximum Likelihood (ML) phylogenetic analysis was conducted as described previously (Henningsen et al. 2024a) on a total of 380 isolates including the 28 newly sequenced isolates from 2024 and a reference panel of 352 previously sequenced isolates collected in Australia (2020 to 2023 collections), the United States of America (USA), South Africa, and Taiwan (Henningsen et al. 2024a; Hewitt et al. 2024; Ho et al. 2025; Miller et al. 2020) (**Figure 2**). The inclusion of *Pca* datasets from other continents allowed a comparative genomic framework to place the 2024 *Pca* isolates within both national and global contexts (Henningsen et al., 2024a). As noted by Henningsen et al. (2024a) the Australian *Pca* collection presents a drastic differentiation from other countries and this pattern was also observed in the samples collected in 2024. Most *Pca* isolates from 2024 clustered within the previously established genotypic lineages (L1 to L18). A novel lineage, designated as L19, was identified from a single isolate (24WA15) collected from WA (**Table 1; Figure 1A,B; Figure 3**). The isolate 24WA15 formed a distinct branch outside of the 18 previously defined lineages (L1–L18), with high bootstrap support (≥ 80%) and a relatively long branch length. The topology of the ML phylogenetic tree indicates that L19 falls within the broader group of Australian lineages (L4 to L17) that from a locally recombining population derived from the L3 and L4 founder lineages (Henningsen et al. 2024a) and is most similar to L7, L8, and L10, all which are present in WA. Thus, we anticipate L19 represents an additional recombinant lineage and unlikely to be an exotic introduction into the continent.

**Fig 2.**
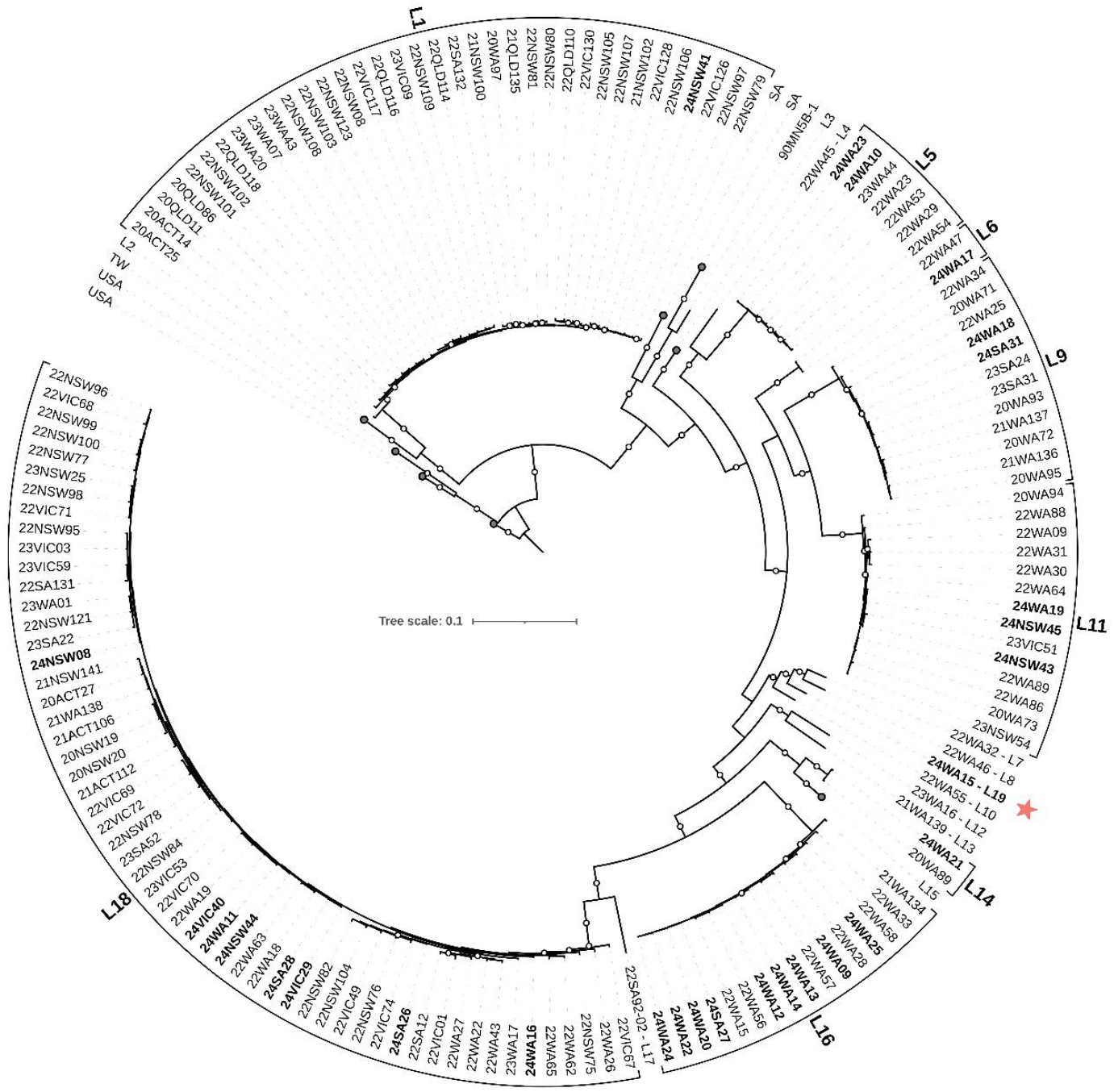
A Maximum likelihood phylogenetic tree of 380 *Pca* isolates worldwide. A midpoint-rooted maximum likelihood tree was constructed using 396,452 biallelic SNPs, with 500 bootstrap replicates. Bootstrap support values ≥ 80% are indicated by white circles. Isolates collected in 2024 are shown with bold text. Previously characterised Australian lineages (L1– L18) are shown with brackets. The newly described L19 is shown with a pink coloured star. Scale bar represents mean substitutions per site.

**Fig. 3.**
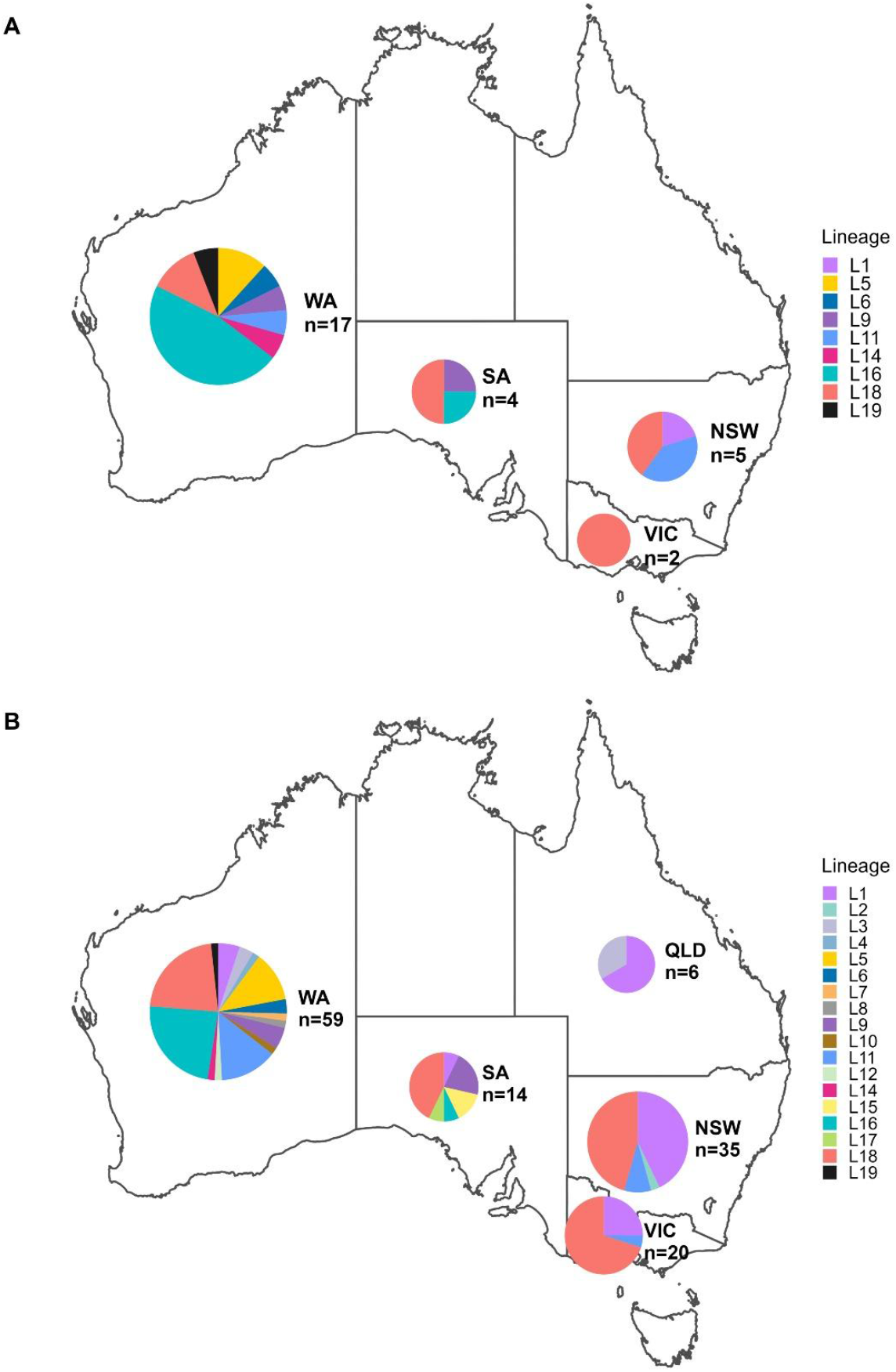
Distribution and number of *Pca* isolates and lineages across Australia. **A**. Collection of *Pca* in 2024. **B**. Collection of *Pca* including samples from 2022 to 2024. Pie chart sizes reflect a logarithmic transformation of the total number of *Pca* isolates collected in their respective states. The pie chart for NSW includes isolates from both regions. Colors reflect genotypic lineage assignments.

We also assessed the distribution and frequency of genotypic lineages across Australia in the 2024 season and compared to the complete *Pca* collection (2022-2024) (**Figure 3A,B**). As illustrated, L18 was found in all surveyed regions and accounted for 29% (8 of 28 total) of samples (**Figure 1A,B; Figure 3**), consistent with our previous report of its dominant presence in both 2022 and 2023 (Henningsen et al. 2024a). Although L1 has previously been reported across all regions, in 2024 L1 was only sampled in NSW. We also note the representation of L18 and L11 in NSW and only L18 in VIC. We only detected three genotypic lineages in South Australia (L9, L16 and L18). WA harbors most of the genetic diversity, with eight lineages being sampled in 2024. Although L16 was not reported 2023 after being detected in 2022, the collection of 2024 showed its prominent presence, particularly in WA, where it accounted for approximately one-third (9 of 28 total, and 8 of 17 from WA) of the sequenced isolates. The detection of L19 adds further evidence that the *Pca* population in WA harbors extensive genetic diversity that we have not yet fully captured.

To complement SNP-based phylogenetic inference, we additionally employed *k-*mer containment analysis as an independent approach to assess genome-wide sequence identity (Henningsen et al. 2024a; Sperschneider et al. 2023) to the Australian and USA pangenome collection of haplotype references (Henningsen et al. 2024a). Briefly, thresholds for haplotype containment were defined as ≥ 99.98 % *k*-mer identity and ≥ 99.65 % shared *k*-mers (**Supplemental Table 3**). This analysis confirmed the presence of the appropriate haplotypes in the 24 isolates assigned to lineages L1, L9, L11, L14, L16 and L18, and showed that L19 does not contain any of the previously defined nuclear haplotypes present in the current pangenome (Henningsen et al. 2024a). Thus, future expansion of the *Pca* pangenome should therefore include 24WA15 to closely examine its precise origin through haplotype assembly, recombination block analysis and ancestry signals.

Pathogenicity assays showed that 24WA15 (L19) is virulent on oat lines Pc40, Pc45, Pc46, Pc54, Pc60, Pc62, Pc64, Pc67, Pc70 and IA B605Xsel in addition to the susceptible control oat lines Marvelous and Swan (**Figure 1A; Table 1**). Like the other genotypic lineages present in WA (e.g., L14), the virulence profile of 24WA15 is limited to a relatively small number of *Pc* genes and contrasts with the broadly virulent lineage L1 common in the Eastern side of the country but detected rarely in WA (Henningsen et al. 2024a) (Figure 1A; Figure 3A). To further explore the industry risk of L19 we assessed the virulence profile of 24WA15 and compared it to samples from L1, L2, L3, L17, and L18 across a small set of commercial oat cultivars including the cultivar Swan as a susceptible control (Supplemental Figure 2). Notably, all commercial cultivars were susceptible to L1, and most varieties were resistant to L2 and L17. Lineages L18 and L19 were similar in virulence.

Comparison of the genotypic and phenotypic data (Figure 1A) also highlights the role of mutation in generating virulence diversity within genetic groups. Isolates within the same genetic lineage differ in virulence on some differential lines likely due to stepwise mutation events in specific avirulence gene loci within these clonal lineages.

While most *Pca* samples included in this study were collected from wild oats, we and other have previously shown that there is no distinction between *Pca* populations found on wild or cultivated oats (Henningsen et al. 2024a; Hewitt et al. 2024; Park et al. 2025). Wild oats are extremely prevalent in Australia and are found near the edges of oat plantations, so this collection represents the disease challenges present in cultivated oat fields. Whilst the 2024 *Pca* collection reveals some changes in pathogen diversity both at the phenotypic and genotypic level, we note the sampling limitations and biases that should be considered before concluding this is due to shifts in host-related selection pressures. The reported genetic recombination of *Pca* in WA (Henningsen et al. 2024a) indicates involvement of a small number of less virulent parental genotypes (i.e., L3 and L18) which has not led to a substantial impact on the emergence of new virulent races. Nevertheless, the potential for recombination to expand to additional lineages, especially the highly virulent L1 lineage, poses a risk for future emergence of novel virulence combinations. Hence it is important to keep monitoring for new lineages in the Australian *Pca* population. Future research will determine whether L19 is a short-lived variant or has the capacity to persist and spread across regions. However, the detection of L19 as part of this relatively small collection warrants increased focus in surveillance, particularly in WA.

Together, our findings highlight the value of using the *Pca* haplotype pangenome as a resource to enhance pathogen monitoring such as in the case of the 2024 *Pca* survey. This study demonstrates the power of coupling phenotypic and genotypic data in advancing our understanding of the evolution of *Pca* population in Australia. Future surveillance efforts, particularly in WA, will be key to assess the completeness of the current assignment of genotypic lineages and support breeding programs.

## Data availability

Genomic short reads are available on the CSIRO Data Access Portal at https://data.csiro.au/collection/csiro:69616. Data underlying summary statistics and scripts used to perform analyses and generate figures are available on GitHub https://github.com/Renee-SU/Pathotyping_2024.

## Acknowledgments

We thank Allan Rattey (InterGrain), Kylie Chambers (DPIRD, WA), Andrea Hills (DPIRD, WA), Jason Bradley (DPIRD, WA), Joel Kidd (DPIRD, WA), Brad Baxter (New South Wales Department of Primary Industries), Michael Ayliffe (CSIRO, ACT) for contributing samples to the 2024 collection used in this study.

## Supplemental Information

**Supplemental Figure 1.**
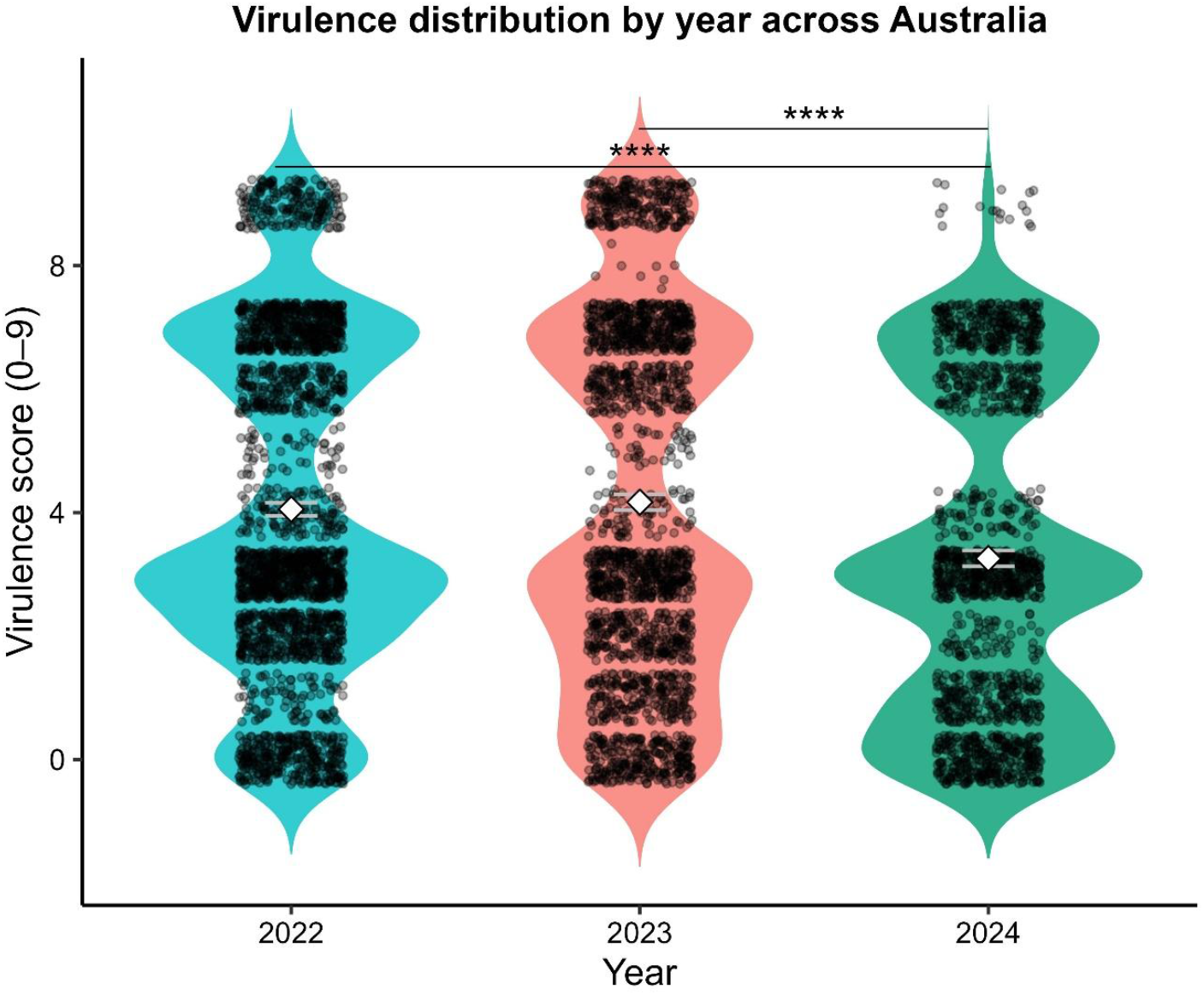
Distribution of infection scores of *Pca* isolates by year against oat differential lines. A box plot was drawn inside each violin plot to indicate median and inter-quartile range values. *p*-value was calculated using the Wilcoxon rank-sum test. **** indicates a highly significant difference (p < 0.001).

**Supplemental Figure 2.**
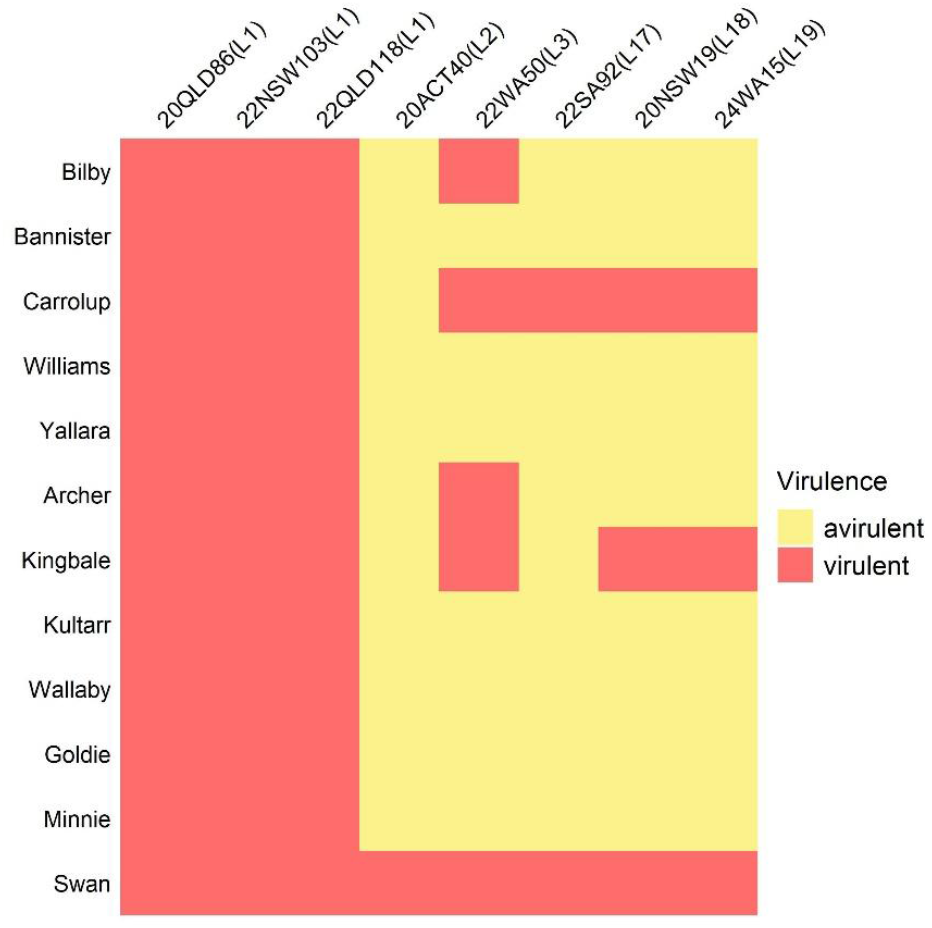
Heatmap showing virulence profiles of *Pca* isolates representing L1, L2, L3, L17, L18, L19 (x-axis) on 11 oat commercial cultivars, plus Swan as susceptible control (y-axis). High infection scores indicating high virulence (susceptibility) are shown in red, and lower infection scores indicating avirulence (resistance) are shown in yellow.

**Supplemental Table 1**. Virulence scores of each *Pca* isolate collected in 2024 across the oat differential set.

**Supplemental Table 2**. Isolate metadata and sequencing information produced in this study.

**Supplemental Table 3**. Haplotype *k*-mer containment from *Pca* Illumina data.

## Literature Cited

Bureau of Meteorology. 2025. Recent and historical rain maps. https://www.bom.gov.au/climate/maps/rainfall [accessed 26 September 2025].

Carson, M.L. 2011. Virulence in oat crown rust (Puccinia coronata f. sp. avenae) in the United States from 2006 through 2009. Plant Dis. 95:1528–1534. 10.1094/PDIS-09-10-0639

Chong, J., Leonard, K. J., and Salmeron, J. J. 2000. A North American system of nomenclature for Puccinia coronata f. sp. avenae. Plant Dis. 84:580–585. 10.1094/PDIS.2000.84.5.580

Danecek, P., Auton, A., Abecasis, G., Albers, C.A., Banks, E., DePristo, M.A., Handsaker, R.E., Lunter, G., Marth, G.T., Sherry, S.T. and McVean, G. 2011. The variant call format and VCFtools. Bioinformatics. 27(15):2156–2158. 10.1093/bioinformatics/btr330

Figueroa, M., Dodds, P.N., and Henningsen, E.C. 2020. Evolution of virulence in rust fungi— multiple solutions to one problem. Curr. Opin. Plant Biol. 56:20–27. 10.1016/j.pbi.2020.02.007

Henningsen, E.C., Lewis, D., Nazareno, E., Huang, Y-F., Steffenson, B. J., Boesen, B., Kianian, S.F., Stone, E., Dodds, P.N., Sperschneider, J., and Figueroa, M. 2024a. A high-resolution haplotype collection uncovers somatic hybridization, recombination and intercontinental movement in oat crown rust. PLoS Genet. 20 (11):e1011493 10.1101/2024.03.27.583983

Henningsen, E.C., Lewis, D., Nguyen, D.T., Sperschneider, J., Kianian, S.F., Stone, E., Dodds, P.N., and Figueroa, M. 2024b. Virulence patterns of oat crown rust in Australia - Season 2022. Plant Dis. 108(7):1959–1963. 10.1094/PDIS-09-23-1973-SC

Hewitt, T.C., Henningsen, E.C., Pereira, D., McElroy, K., Nazareno, E.S., Dugyala, S., Nguyen-Phuc, H., Li, F., Miller, M.E., Visser, B. and Pretorius, Z.A, Boshoff, W.H., Sperschneider, J., Stukenbrock, E.H., Kianian, S.F., Dodds, P.N., and Figueroa, M. 2024. Genome-enabled analysis of population dynamics and virulence-associated loci in the oat crown rust fungus Puccinia coronata f. sp. avenae. Mol Plant Microbe In. 37:290–303. 10.1094/MPMI-09-23-0126-FI

Ho, C.Y., Henningsen, E.C., Chen, S.T., Ariyawansa, H.A., Nazareno, E.S., Sperschneider, J., Dodds, P.N., Riddle, J., Kianian, S.F., Figueroa, M., and Huang, Y.F. 2025. Confirmation of oat crown rust disease in Taiwan. Plant Dis. 109(3): 691–697. 10.1094/PDIS-04-24-0838-RE

McCallum, B.D., Fetch, T., and Chong, J. 2007. Cereal rust control in Canada. Aust. J. Agric. Res. 58(6):639–647. 10.1071/AR06145

McDonald, B.A., and Linde, C. 2002. The population genetics of plant pathogens and breeding strategies for durable resistance. Euphytica, 124(2): 163–180. 10.1023/A:1015678432355

Miller, M.E., Nazareno, E.S., Rottschaefer, S.M., Riddle, J., Dos Santos Pereira, D., Li, F., Nguyen-Phuc, H., Henningsen, E.C., Persoons, A., Saunders, D.G.O., Stukenbrock, E., Dodds, P.N., Kianian, S.F., and Figueroa, M. 2020. Increased virulence of Puccinia coronata f. sp. avenae populations through allele frequency changes at multiple putative Avr loci. PLoS Genet. 16:e1009291. 10.1371/journal.pgen.1009291

Moreau, E.L., Riddle, J.M., Nazareno, E.S., and Kianian, S.F. 2024. Three decades of rust surveys in the United States reveal drastic virulence changes in oat crown rust. Plant Dis. 108(5):1298–1307. 10.1094/PDIS-09-23-1956-RE

Nazareno, E.S., Li, F., Smith, M., Park, R.F., Kianian, S.F., and Figueroa, M. 2018. Puccinia coronata f. sp. avenae: a threat to global oat production. Mol. Plant Pathol. 19(5):1047–1060. 10.1111/mpp.12608

Nguyen, D.T., Henningsen, E.C., Lewis, D., Mago, R., Sperschneider, J., Stone, E.A., Dodds, P.N., and Figueroa, M. 2025. Characterisation of virulence of Puccinia coronata f. sp. avenae in Australia in the 2023 growing season. Plant Dis. 109(8):1625–1629. 10.1094/PDIS-09-24-2049-SC

Park, R.F., Kosman, E., Roake, J., Ding, Y., Winter, B., Zwer, P.K., and Singh, D. 2025. Long-Term Studies of Puccinia coronata f. sp. avenae in Australia Reveal High Pathogenic Diversity, Regional Virulence Differences, Evidence of Clonality and Rapid Emergence of Virulence Matching Deployed Host Resistance. Plant Pathol. 74 (6):1778–1795. 10.1111/ppa.14125

Simons, M.D. 1985. Crown rust. Pages 142–172 in: The Cereal Rusts Vol II: Diseases, distribution, epidemiology and control. A. P. Roelfs and W. R. Bushnell eds., Academic Press, Orlando, FL. 10.1016/C2013-0-10449-7

Sperschneider, J., Hewitt, T., Lewis, D.C., Periyannan, S., Milgate, A.W., Hickey, L.T., Mago, R., Dodds, P.N., and Figueroa, M. 2023. Nuclear exchange generates population diversity in the wheat leaf rust pathogen Puccinia triticina. Nat. Microbiol. 8(11): 2130–2141. 10.1038/s41564-023-01494-9

USDA-ARS CDL. 2014. Oat Loss to Rust. USDA-ARS Cereal Disease Laboratory. https://www.ars.usda.gov/ARSUserFiles/50620500/Smallgrainlossesduetorust/2014loss/2014oatloss.pdf [accessed 21 September 2025].

